# Polymetallic nodules are essential for food-web integrity of a prospective deep-seabed mining area in Pacific abyssal plains

**DOI:** 10.1101/2021.02.11.430718

**Authors:** Tanja Stratmann, Karline Soetaert, Daniel Kersken, Dick van Oevelen

**Affiliations:** Department of Estuarine and Delta Systems, NIOZ Royal Netherlands Institute for Sea Research, P.O. Box 140, 4400 AC Yerseke, The Netherlands; Department of Earth Sciences, Utrecht University, Vening Meineszgebouw A, Princetonlaan 8a, 3584 CB Utrecht, The Netherlands; HGF MPG Joint Research Group for Deep-Sea Ecology and Technology, Max Planck Institute for Marine Microbiology, Celsiusstraße 1, 28359 Bremen, Germany; German Centre for Marine Biodiversity Research (DZMB), Senckenberg am Meer, Südstrand 44, 26382 Wilhelmshaven, Germany; Marine Zoology, Senckenberg Research Institute and Nature Museum, Senckenberganlage 25, 60325 Frankfurt am Main, Germany

**Keywords:** binary food web, DISCOL, anthropogenic impact, interaction matrix

## Abstract

Polymetallic nodule fields provide hard substrate for sessile organisms on the abyssal seafloor between 3,000 and 6,000 m water depth. Deep-seabed mining targets these mineral-rich nodules and will likely modify the consumer-resource (trophic) and substrate-providing (non-trophic) interactions within the abyssal food web. However, the importance of nodules and their associated sessile fauna in supporting food-web integrity remains unclear. Here, we use seafloor imagery and published literature to develop highly-resolved trophic and non-trophic interaction webs for the Clarion-Clipperton Fracture Zone (CCZ, central Pacific Ocean) and the Peru Basin (PB, south-east Pacific Ocean) and to assess how nodule removal will modify these networks. The CCZ interaction web included 1,028 compartments connected with 59,793 links and the PB interaction web consisted of 342 compartments and 8,044 links. We show that knock-down effects of nodule removal resulted in a 17.9% (CCZ) to 20.8% (PB) loss of all compartments and 22.8% (PB) to 30.6% (CCZ) loss of network links. Subsequent analysis identified stalked glass sponges living attached to the nodules as key structural species that supported a high diversity of associated fauna. We conclude that polymetallic nodules are critical for food-web integrity and that their absence will likely result in reduced local benthic biodiversity.

## 1. Introduction

Abyssal plains, most seafloor between 3,000 and 6,000 m water depth, have been relatively untouched by anthropogenic impacts due to their extreme depths and distance from continents (Ramírez-Llodrà et al., 2011). This may change in the near future with prospected deep-seabed mining, a commercial activity that will extract polymetallic nodules from the seabed for their metal content (Hein & Koschinsky, 2014). Polymetallic nodules are slow-growing (10s mm My^-1^, (Hein, 2016)) precipitates of predominantly manganese oxides and iron oxy-hydroxides that contain metals such as nickel, cobalt, and copper (Hein & Koschinsky, 2014). They are found partially buried on the sediment surface with a mean nodule wet weight of 4.5 kg m^-2^ (central Indian Ocean Basin) to 15 kg m^-2^ (Clarion-Clipperton Fracture Zone, CCZ, North-East Pacific) (Kuhn et al., 2017). Nodules provide the rare commodity of hard substrate in the abyss that is used by sessile fauna, e.g., Porifera, Antipatharia, Alcyonacea, or Ascidiacea, and by mobile fauna, e.g., Cephalopoda (Amon et al., 2016; Purser et al., 2016; Vanreusel et al., 2016). In this way, nodules add a network of non-trophic interactions (i.e., non-consumptive relationships between taxa) among sessile organisms and their associated fauna, the so-called epibiota, to the classical trophic interactions that are known for abyssal food webs.

Abyssal food-web studies traditionally focus on trophic interactions (i.e., consumer-resource relationship; e.g., (Aberle & Witte, 2003; Dunlop et al., 2016; Iken et al., 2001; de Jonge et al., 2020; van Oevelen et al., 2012; Sweetman & Witte, 2008)), because life at abyssal plains is strongly food-limited (Smith et al., 2008). The top sediment layer is typically well-oxygenated, reducing the importance of habitat modification by sediment ventilation or bioturbating fauna (Meysman et al., 2006). Deep-seabed mining will remove polymetallic nodules from the seafloor and thereby modify the network of non-trophic interactions that is an essential part of the integrity of a nodule field food web on the abyssal seabed. It is therefore of imminent importance to understand how polymetallic nodules support trophic and non-trophic interactions in the abyss.

Here, we develop high-resolution trophic and non-trophic interaction webs for two nodule-rich areas in the Pacific Ocean, i.e., the CCZ and the Peru Basin in the South-East Pacific, based on an extensive literature compilation, supplemented with the analysis of high-resolution seafloor images. An important element of these webs is that they include trophic and non-trophic dependencies between taxa and between polymetallic nodules and taxa. Deep-seabed mining impacts are subsequently simulated by assessing the knock-down effects through the food web following the initial removal of only the non-trophic links among nodules and taxa. We focused our analysis on the following questions:

1. Can the importance of polymetallic nodules as hard substrate in abyssal plains be quantified?
2. Are faunal taxa that provide habitat structure (i.e., structural faunal taxa) more important for food-web integrity than highly connected faunal taxa?

## 2. Materials and Methods

### 2.1 Study sites

The CCZ (0°N, 160°W to 23.5°N, 115°W (International Seabed Authority, 2011)) is bound by the Clarion Fracture Zone in the north and the Clipperton Fraction Zone in the south. Water depth ranges from 3,900 m in the east (Purser et al., 2016) to 5,300 m in the west (Jung et al., 1998) with an average water depth of 4,794 m (Wedding et al., 2013). The CCZ is part of the ‘mesotrophic’ abyss (Hannides & Smith, 2003) and experiences a gradient in particulate organic carbon (POC) input from north (1.3 mg organic C m^-2^ d^-1^) to south (1.8 mg organic C m^-2^ d^-1^) (Vanreusel et al., 2016). The sediment is largely siliceous, silty clay (International Seabed Authority, 2010) and contains nodules at a density between 0 and ~30 kg m^-2^ with a mean density of 15 kg m^-2^ (Kuhn et al., 2017) and a mean diameter of 3.6 cm (Schoening et al., 2017). Because of this high nodule density, the CCZ is an important area for future nodule mining operations and (until September 2020) the International Seabed Authority has granted 18 15-year contracts for the exploration of polymetallic nodules in the CCZ.

The Peru Basin is located in the southern Pacific (5°S to 24°S) and extends from the Atacama Trench west of Peru westwards (to 110°W) into the Pacific (Bharatdwaj, 2006; Klein, 1993). The water depth ranges from 3,900 to 4,300 m (Glasby, 2006) and sedimentation rates are between 0.4 and 2.0 cm kyr^-1^ (Haeckel et al., 2001). Surface sediments have an organic carbon content of 0.5 to 1.0 weight % and the oxygen concentration decreases from 132 μmol L^-1^ at the sediment-water interface to 0 μmol L^-1^ at 6 to 15 cm sediment depth (Haeckel et al., 2001). Mean nodule density in the Peru Basin is >10 kg m^-2^ (Glasby, 2006) and mean nodule diameter is 7.8 cm (Schoening et al., 2017); industrial polymetallic nodule extraction, however, is not anticipated.

### 2.2 Data compilation

Detailed food webs were developed for the size classes protozoan and metazoan meiobenthos, macrobenthos, invertebrate megabenthos, and fish, while explicitly taking into account whether each taxon lives in the sediment, is associated with other taxa, or is attached to polymetallic nodules. For this purpose, a systematic literature review was conducted to identify all taxa (if possible on genus level) of the study areas following the PRISMA guidelines for systematic reviews and meta-analyses (Moher et al., 2009). The key words ‘polymetallic nodules’, ‘manganese nodule fauna’, ‘manganese nodule’, ‘Clarion-Clipperton Fracture Zone’, ‘Peru Basin’, ‘DISCOL’ identified 913 publications in the database ‘Web of Knowledge’ (Fig. S1). Additional 113 records were identified through other sources, 121 publications were listed on the ‘DISCOL’ homepage (https://www.discol.de/literatur), and 66 articles were published in three special issues in *Deep-Sea Research Part II: Topical Studies in Oceanography* (Thiel, 2001), *Frontiers in Marine Sciences* (Santos et al., 2019), and *Biogeosciences* (Middelburg et al., 2018). After duplicates were removed, title and abstract of 854 records were screened for relevancy and 671 records were excluded because they did not report benthos from the CCZ or the Peru Basin. Subsequently, a total of 183 full-text records were assessed for eligibility. The full text was unavailable for three records that were therefore left out. Another 24 publications were excluded as they did not report benthic taxa from the CCZ or the Peru Basin and eight studies were omitted because they presented meta-studies or reviews and not primary research. In the end, 148 studies or datasets were used to develop the trophic and non-trophic interaction matrices. All references can be found in Table S2.

The extensive literature survey was supplemented by two novel datasets of high-resolution seafloor images (Fig. S1). During RV *Sonne* cruise SO242-2 to the Peru Basin (Boetius, 2015) and during RV *Sonne* cruise SO239 to the CCZ (Martínez Arbizu & Haeckel, 2015), 1,544 seafloor images were taken with the remotely operated vehicle ‘ROV Kiel 6000’ (Geomar, Kiel, Germany) and Porifera and its associated fauna were identified.

All metazoan and protozoan specimens, that were discovered in the qualitative synthesis and the picture analysis, were compiled in a database. For each specimen, the taxonomic ranks phylum, class, order, family, and genus were determined using the database ‘World Register of Marine Species: WoRMS’ (Horton et al., 2018) and the genus level of all taxa was selected as food-web compartment. However, when a specimen could not be classified at this taxonomic rank, the lowest taxonomic rank above genus level, to which it could be identified, was chosen as food-web compartment (Table S3). Recorded specimens were furthermore classified as meiobenthos (>32 μm; (Ahnert & Schriever, 2001; Radziejewska, 2002)), macrobenthos (>250 μm/ >500 μm; (Borowski, 2001; Borowski & Thiel, 1998)), invertebrate megabenthos (>1 cm), and fish based on information from original publications or based on information published in “WoRMS” (Horton et al., 2018). It was also noted whether organisms were reported or observed attached to a polymetallic nodule, sitting on or living in sediment, or were associated with an organism attached to a nodule.

### 2.3 Trophic interaction matrix

Feeding preferences/ prey types of each specimen in the dataset were identified by inquiries of published literature and the database ‘WoRMS’ (Horton et al., 2018) (Data file S1 and S2). One binary trophic interaction matrix per location was developed by connecting all food-web compartments via all reported trophic links to its prey/ food source. Hence, when a specific fish genus preferably feeds on crustaceans, this fish genus compartment was connected via trophic links to all compartments in the food web that include crustaceans. However, it was assumed that predatory megabenthos would not feed on meiobenthos unless they are known to catch their prey by digging in the seafloor.

Bacteria, carrion, dissolved organic matter (DOM), fungi, (labile) sedimentary detritus, phytodetritus, Protozoa, Rotifera, and particulate organic matter (POM) suspended in the water column were reported in the literature as food sources for abyssal food webs and were consequently included (Data file S1 and S2).

### 2.4 Non-trophic interaction matrix

The binary non-trophic interaction matrix contained all non-trophic links between food-web compartments and the additional compartment ‘polymetallic nodules’. Commensal relations among faunal compartments that were reported in the database (Data compilation section) were implemented in the matrix as present or absent. All faunal compartments, for which the literature or the image analysis indicated that they would use nodules as substrate, were linked to the nodule compartment. Whenever the taxon of a faunal compartment was observed attached to a nodule and living on/ in the sediment or associated with a faunal compartment and freely, the present non-trophic relation was defined as ‘facultative’. In contrast, when the taxon occurred exclusively attached to nodules or only in association with specific faunal compartments, the present non-trophic relation was defined as ‘obligatory’.

### 2.5 Network indices

The trophic interaction matrix was used to calculate the well-established network indices: ‘number of interaction web compartments’ S, ‘number of network links’ L, ‘link density’ D (*D*=*L/S*), and ‘connectance’ C (*C*=*L/S2*), i.e., the fraction of realized links compared to all possible links (Pimm et al., 1991).

### 2.6 Assessment of the absence of nodules and the absence of specific faunal compartments

The effect of polymetallic nodules on the food-web structure was investigated by removing the nodule compartment from the non-trophic interaction matrix and assessing the resulting knock-down on other compartments (i.e., the secondary extinction of other compartments because of the loss of the nodule compartment). Hence, all compartments directly depending on the nodules were removed first. In a subsequent iteration, all compartments depending on a removed compartment through a non-trophic or exclusively trophic (i.e., when preying only on the removed compartment) interaction were removed, and so on. After all lost compartments were identified, the previously collected information about the type of non-trophic interaction (facultative vs. obligatory) was used to identify whether the loss was a result of facultative or obligatory dependence.

To compare the loss of non-trophic links through nodule absence against the removal of trophic links, a comparable knock-down analysis of the ‘most connected taxa’ and the ‘taxa with most non-trophic interactions’ was also carried out. The most connected taxon is defined as the food-web compartment with most trophic links, *sensu* van der Zee et al. (van der Zee et al., 2016). The ‘highest impact taxon’ (van der Zee et al., 2016) was identified by removing each faunal compartment individually and calculating the food-web properties. The compartment whose individual removal resulted in the largest absolute difference in food-web properties between the default food web and the modified food web was defined as said ‘highest impact taxon’.

## 3 Results

### 3.1 Food web of the abyssal plains in the Clarion-Clipperton Fracture Zone

Based on the literature survey and image analysis, we identified 1,018 faunal food-web compartments (Table 1) across the size ranges of protozoan and metazoan meiobenthos (36%), macrobenthos (37%), invertebrate megabenthos (25%), and fish (2%). From the literature survey we further identified feeding preferences for all compartments and we showed that 56% of all faunal compartments were exclusive deposit feeders (265 compartments), filter/suspension feeders (112 compartments), or carnivores (188 compartments) (Fig. 1A). Feeding interactions among nine food-source compartments (bacteria, carrion, dissolved organic matter, fungi, (labile-) sedimentary detritus, phytodetritus, Protoza, suspended detritus) and all faunal compartments resulted in the highest-resolved trophic interaction deep-sea food web to date of 59,793 links (Fig. 2, Table 1). The non-trophic interaction web contained a total of 386 links (Fig. 2). The megabenthic *Hymenaster* sp. (Echinodermata) was the so-called ‘most connected taxon’, i.e. the taxon with most trophic interactions (*sensu* (van der Zee et al., 2016)). The macrobenthic *Abyssarya* sp. (Annelida) was the taxon with most non-trophic (commensal) interactions.

**Figure 1.**
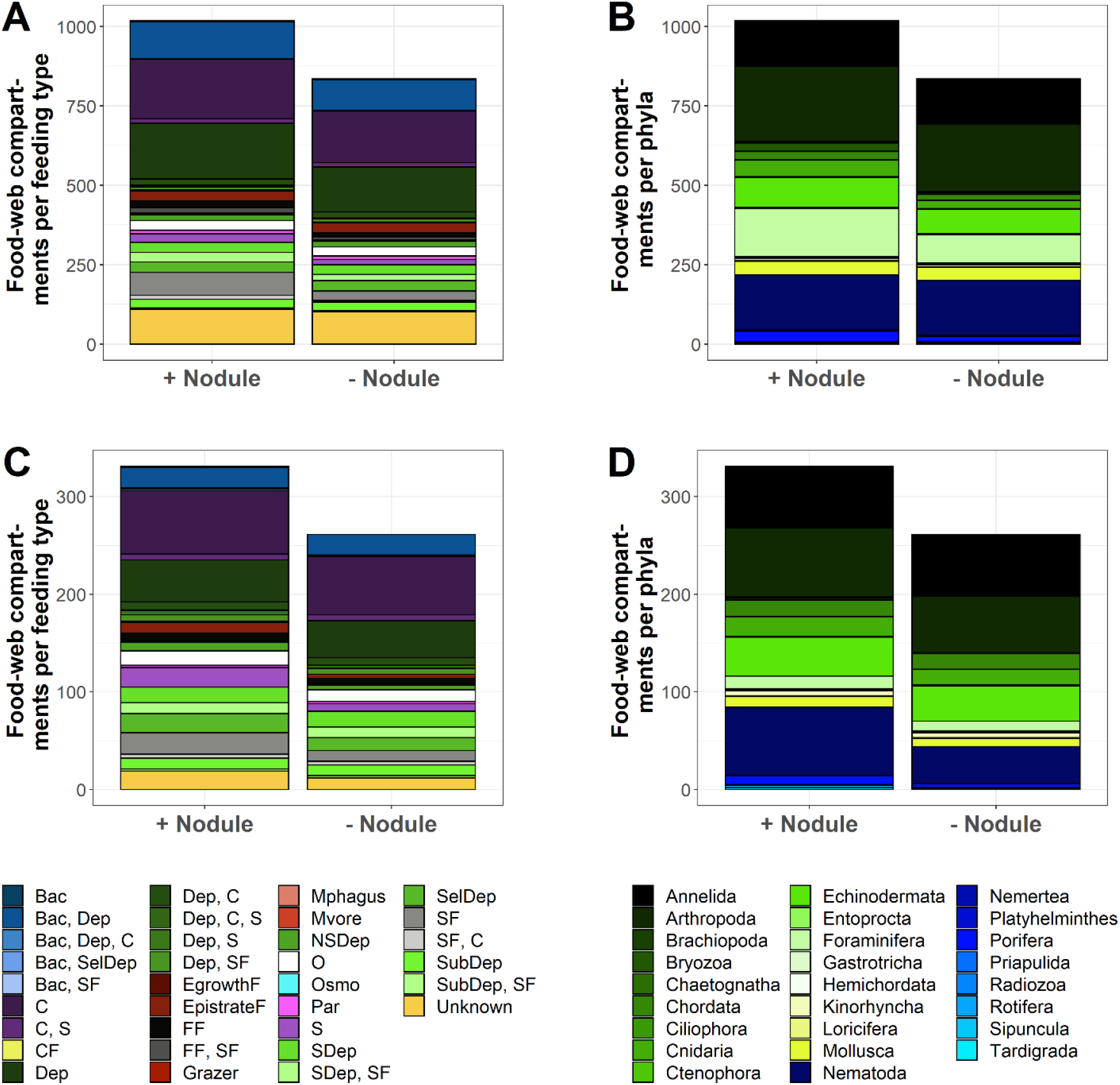
Number of feeding types and phyla present in the Clarion-Clipperton Fracture Zone (CCZ) and the Peru Basin. (A) All compartments that are part of trophic and non-trophic interaction webs of the Clarion-Clipperton Fracture Zone (CCZ) are split into feeding types and shown in the presence (+ nodule) and absence (- nodule) of polymetallic nodules. (B) All compartments from the CCZ interaction webs are divided into phyla and presented in the presence (+ nodule) and absence (- nodule) of nodules. (C) All compartments of the Peru Basin interaction web shown as feeding types in the presence (+ nodule) and absence (- nodule) of nodules. (D) All phyla that occur in the Peru Basin in the presence (+ nodule) and absence (- nodule) of nodules. Abbreviation: Bac = bacterivore, C = carnivore, CF = ciliate feeder, Dep = deposit feeder, EgrowthF = epigrowth feeder, EpistrateF = epistrate feeder, FF = filter feeder, Mphagus = microphagous, Mvore = microvore, NSDep = non-selective deposit feeder, O = omnivore, Osmo = osmotroph, Par = parasitic, S = scavenger, SDep = surface deposit feeder, SelDep = selective deposit feeder, SF = suspension feeder, SubDep =subsurface deposit feeder, Unknown = unknown feeding type.

**Figure 2.**
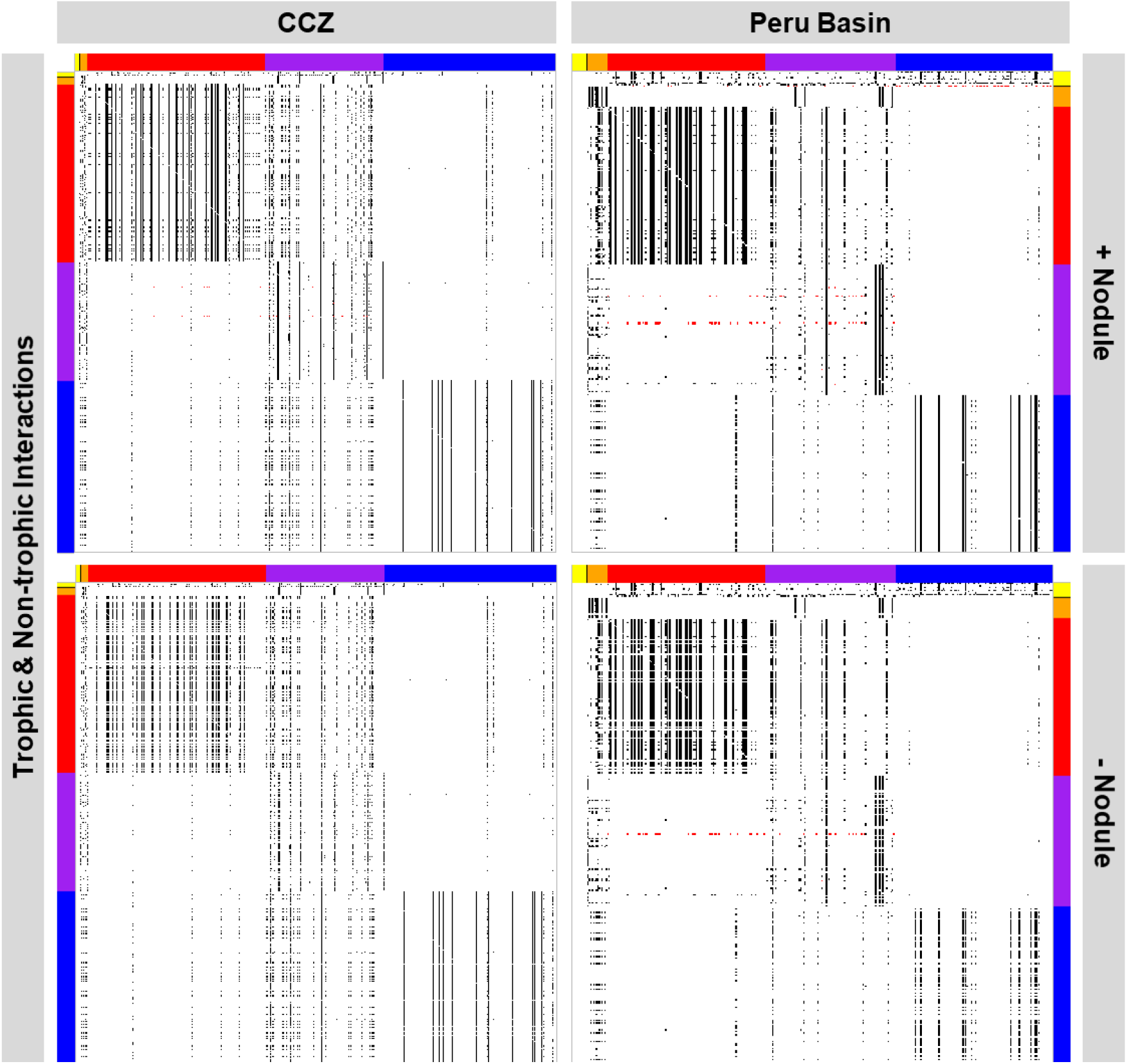
Network links between food-web compartments. Each individual dot represents an interaction between polymetallic nodules (black section) and faunal food-web compartments (orange, red, purple, and blue sections), two faunal food-web compartments, or a faunal food-web compartment and the food-source compartment (yellow section). (Top left) Trophic (black) and non-trophic (red) interactions or links in the Clarion-Clipperton Fracture Zone (CCZ) when polymetallic nodules are present. (Bottom left) All trophic (black) and non-trophic (red) interactions or links that remain in the Clarion-Clipperton Fracture Zone interaction webs after polymetallic nodule removal. (Top right) All trophic (black) and non-trophic (red) interactions or links that exist in the Peru Basin when polymetallic nodules are present. (Bottom right) All trophic (black) and non-trophic (red) interactions or links that remain after polymetallic nodule removals in the Peru Basin interaction webs. Color code of axes: yellow = food sources, black = polymetallic nodules, orange = fish, red = macrobenthos, purple = megabenthos, blue = meiobenthos.

**Table 1.** Changes of network properties dependent on the presence or absence of polymetallic nodules. The network properties were calculated for the trophic interaction webs of the Peru Basin and the Clarion-Clipperton Fracture Zone when polymetallic nodules were present and absent, and without the most connected taxon, without the taxon with most non-trophic links, and without the highest impact taxon. The change in percent with respect to the default web is shown in brackets.

| | Number of interaction web compartments $S$ | Number of network links $L$ | Link density $D$ | Connectance $C$ |
| --- | --- | --- | --- | --- |
| **Clarion-Clipperton Fracture Zone** |  |  |  |  |
| With polymetallic nodules | 1,028* | 59,793 | 58 | $5.70 \times 10^{-2}$ |
| Without polymetallic nodules | 844<br>(-17.9%) | 41,526<br>(-30.6%) | 49<br>(-15.5%) | $5.80 \times 10^{-2}$<br>(+1.75%) |
| Without the most connected taxon *Hymenaster* sp. | 1,027<br>(-9.73 $\times 10^{-2}$) | 58,774<br>(-1.70%) | 57<br>(-1.72 $\times 10^{-2}$) | $5.60 \times 10^{-2}$<br>(-1.75%) |
| Without the taxon with most non-trophic interactions *Abyssarya* sp. | 1,027<br>(-9.73 $\times 10^{-2}$) | 59,316<br>(-0.80%) | 58<br>(0%) | $5.60 \times 10^{-2}$<br>(-1.75%) |
| Without the highest impact taxon *Hyalonema* sp. | 988<br>(-3.89%) | 53,266<br>(-10.9%) | 54<br>(-6.90%) | $5.50 \times 10^{-2}$<br>(-3.51%) |
| **Peru Basin** |  |  |  |  |
| With polymetallic nodules | 342** | 8,044 | 24 | $6.90 \times 10^{-2}$ |
| Without polymetallic nodules | 271<br>(-20.8%) | 6,208<br>(-22.8%) | 23<br>(-4.17%) | $8.50 \times 10^{-2}$<br>(+23.2%) |
| Without the most connected taxon *Actinia* sp. | 341<br>(-0.29%) | 7,795<br>(-3.10%) | 23<br>(-4.17%) | $6.70 \times 10^{-2}$<br>(-2.90%) |
| Without the taxon with most non-trophic interactions *Munidopsis* sp. | 341<br>(-0.29%) | 8,027<br>(-0.21%) | 24<br>(0%) | $6.90 \times 10^{-2}$<br>(0%) |
| Without the highest impact taxon *Caulophacus* sp. | 319<br>(-6.73%) | 7,387<br>(-8.17%) | 23<br>(-4.17%) | $7.30 \times 10^{-2}$<br>(+5.80%) |

\*1,028 interaction web compartments = 1,018 faunal compartments + 9 food source compartments + 1 polymetallic nodule compartment.
\*\*342 interaction web compartments = 331 faunal compartments + 10 food source compartments + 1 polymetallic nodule compartment.

Polymetallic nodule removal led to 18% less faunal food-web compartments (Table 1). Food-web compartments that were lacking consisted of protozoan and metazoan meiobenthos (4%), macrobenthos (50%), invertebrate megabenthos (45%), and fish (0.5%). The removal of nodules caused the reduction of microphagous feeders by 50%, of ‘suspension feeders and carnivores’ by 58%, and of suspension feeders by 58% (Fig. 1A). Furthermore, ‘bacterivore and suspension feeder’ and ‘deposit feeder, carnivore, and scavenger’ disappeared completely (Fig. 1A). The most affected phyla were Bryozoa (81% loss), Cnidaria (52% loss), Platyhelminthes and Porifera (both 50% loss) (Fig. 1B). All compartments that were lost from the food web, except for the fish *Pachycara* sp., were lost due to a loss of non-trophic interactions, i.e., obligatory dependence on nodules as substrate (69%), facultative dependence on nodules (6%), obligatory dependence on other faunal compartments (12%), and the facultative dependence on other organisms (13%) (Table S1). The latter associations between faunal food-web compartments included the commensal relationships with Antipatharia, Alcyonacea, and Pennatulacea (all Cnidaria), Ophiurida (Echinodermata), and Porifera (Table S1).

The taxon with the highest impact, i.e., the faunal compartment whose removal has the largest impact on food-web properties (*sensu* (van der Zee et al., 2016), was the megabenthic hexactinellid sponge *Hyalonema* sp. (Fig. 3A), whose removal resulted in the loss of 4% of the food-web compartments (Table 1). In comparison, the removal of the ‘most connected taxon’ *Abyssarya* sp. (Annelida) did not lead to any knock-down of other faunal compartments (Table 1).

**Figure 3.**
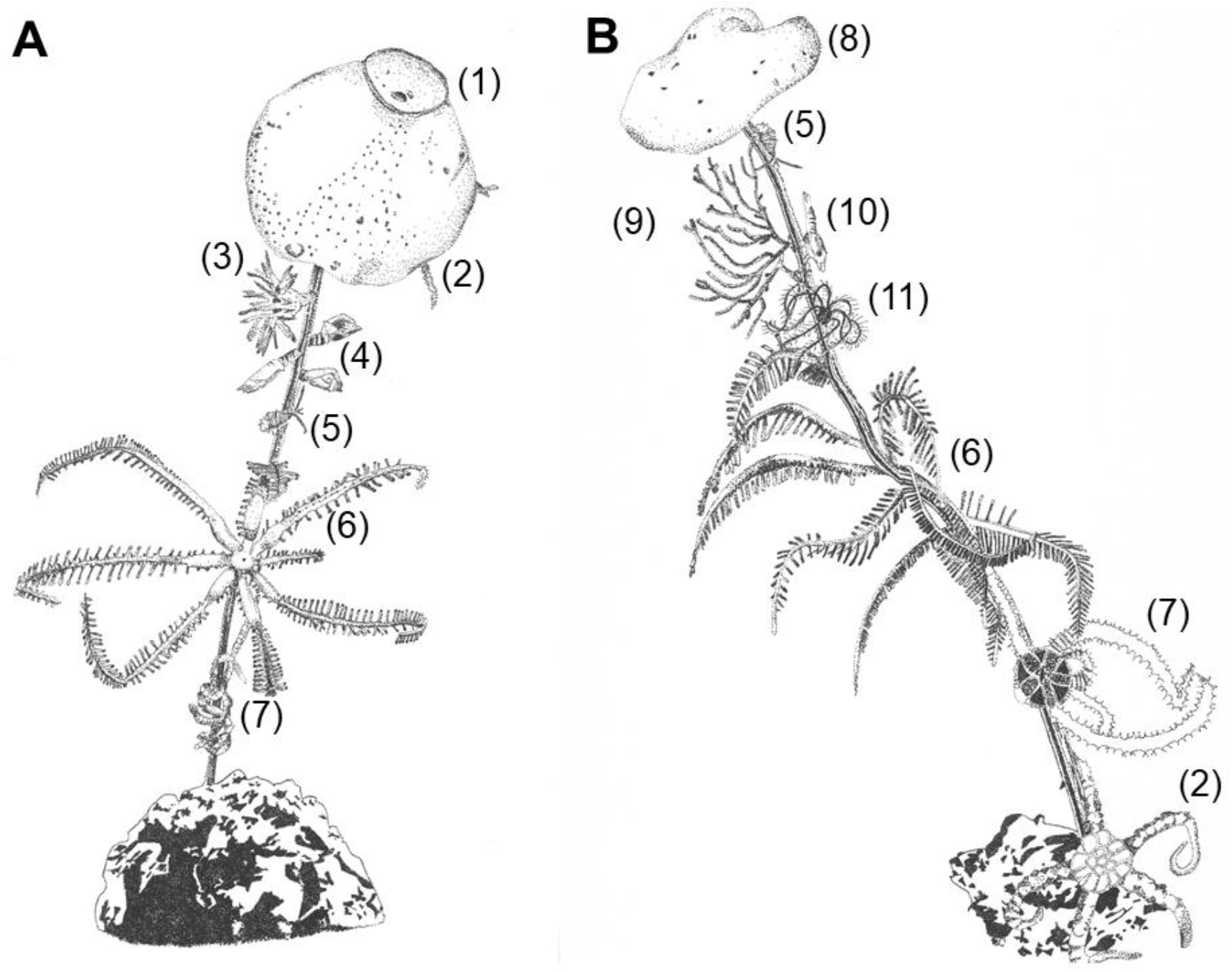
Drawings of highest impact taxa. (A) The highest impact taxon in the Clarion-Clipperton Fracture Zone (CCZ), the hexactinellid sponge *Hyalonema* sp., and (B) the highest impact taxon in the Peru Basin, the hexactinellid sponge *Caulophacus* sp., are shown with their corresponding associated fauna (on Order level) with which they have non-trophic interactions. The taxa presented in the drawings are the following: (1) *Hyalonema* sp., (2) Ophiurida, (3) Actiniaria, (4) Cirripedia, (5) Amphipoda, (6) Brisingida, (7) Ophiacanthida, (8) *Caulophacus* sp., (9) Alcyonacea, (10) Mysida, and (11) Comatulida. Illustrations by Tanja Stratmann.

### 3.2 Food web of the abyssal plains in the Peru Basin

Inquiring the literature and seabed photographs, we found 331 faunal food-web compartments (Table 1), of which 34% belonged to metazoan and protozoan meiobenthos, 34% to macrobenthos, 28% to invertebrate megabenthos, and 4% to fish. Our literature survey indicated that 65% of all faunal compartments were carnivores (65 compartments), deposit feeders (99 compartments), filter/ suspension feeders (29 compartments), and ‘bacterivores and deposit feeders’ (23 compartments) (Fig. 1C). Feeding interactions among ten food-source compartments (bacteria, carrion, dissolved organic matter, fungi, (labile) sedimentary detritus, phytodetritus, suspended detritus, Protozoa, and Rotifera) and all faunal comparments resulted in 8,044 links in the trophic interaction web (Fig. 2, Table 1). The non-trophic interaction web included 215 links (Fig. 2). The megabenthic suspension feeding and carnivorous *Actinia* sp. (Cnidaria) was the most connected taxon and had 244 trophic links. The taxon with most non-trophic (commensal) interactions was the megabenthic *Munidopsis* sp. (Arthropoda).

The removal of nodules resulted in the loss of 21% of all faunal compartments (Table 1), of which 49% belonged to meiobenthos, 24% to macrobenthos, 26% to invertebrate megabenthos, and 1% to fish. When nodules were removed, 50% of all suspension feeders, 60% of all scavengers, and 64% of all epistrate feeders were lost (Fig. 1C). Additionally, all bacterivore, ‘bacterivore, selective deposit feeder’, ‘deposit feeder, carnivore, scavenger’, epigrowth feeder, and microvore were absent (Fig. 1C). Highest losses occurred in Bryozoa (50% loss), Porifera (56% loss), Brachiopoda and Tardigrada (both 100% loss) (Fig. 1D). Except for the fish *Pachycara* sp. that was lost due to the loss of its food source, all other compartments were removed because non-trophic interactions disappeared. Twenty-nine percent of the lost compartments were removed as they obligatory depended on nodules as substrate and 38% because of their facultative dependence on the nodules (i.e., the compartment was not always found attached to nodules) (Table S1). The remaining 33% were removed because of their obligatory (29%) and facultative (4%) commensal relationship with the megabenthic hexactinellid sponges *Bathyxiphus* sp., *Caulophacus* sp., *Hyalonema* sp., and *Hyalostylus* sp. (Table S1).

The taxon with highest impact was the megabenthic hexactinellid sponge *Caulophacus* sp. (Fig. 3B) whose removal resulted in the loss of 23 compartments which is 7% of all faunal compartments (Table 1). In contrast, the removal of the taxon with most trophic interactions *Actinia* sp. (Cnidaria) did not result in any knock-down loss of other food-web compartments (Table 1).

### 3.3 Changes in network properties of both abyssal food webs

Removal of polymetallic nodules as hard substrate resulted in a substantial and consistent reduction in the number of network links for both sites (23% in the Peru Basin and 31% in the CCZ; Fig. 2, Table 1). Link density, i.e., the average number of links per interaction web compartment, was decreased by 4% in the Peru Basin, but by 16% in the CCZ, whereas connectance C, i.e., the fraction of realized links (Hall & Raffaelli, 1993), increased by 23% in the Peru Basin and by 2% in the CCZ (Table 1).

In contrast to nodule removal, the removal of the most connected taxa and the taxa with most non-trophic links decreased the number of network links by only 3% and 0.2%, respectively, in the Peru Basin and by 2% and 0.8%, respectively, in the CCZ (Table 1). In comparison, when the taxa with most non-trophic interactions were removed, the number of network links was 8% (Peru Basin) and 11% (CCZ) lower than the number of links in the corresponding default interaction webs. Link density was not affected by the removal of the taxon with most non-trophic links, but removing the most connected taxon caused a loss in link density of 4% in the Peru Basin and of 7% in the CCZ (Table 1). The connectance decreased slightly when the most connected taxa and the taxa with most non-trophic interactions were removed, but the connectance was unaffected when the taxon with most non-trophic interactions in the CCZ was removed (Table 1). Removing the highest impact taxon resulted in an increase in connectance (+6%) in the Peru Basin, but in a small loss in connectance (−4%) in the CCZ.

## 4 Discussion

### 4.1 Model limitations

The highly resolved binary food webs for the Peru Basin and the CCZ are based on systematic meta-analyses and seabed images, but this comes with inevitable limitations: The literature included in this study did not use consistent lower and upper sieve size limits for meio- and macrobenthos. Furthermore, only few studies reported the use of an upper sieve size for meiobenthos (e.g., (Mahatma, 2009)), whereas the majority of the studies did not report any upper sieve size limit and likely included all infauna that was retained on sieves with the lower size limit. Hence, smaller macrobenthos might be classified as meiobenthos and therefore was erroneously implemented as meiobenthos in the food-web models. In return, many authors used the term ‘meiobenthos’ synonymously to ‘meiofauna’ (e.g., (Giere, 2009)) instead of distinguishing between the statistical entity ‘meiobenthos’ based on body size and the taxonomic category ‘meiofauna’ (McIntyre, 1969). Therefore, when authors (e.g., (Borowski, 2001)) stated that specific taxa were excluded from the analysis of a specific size range because the authors used taxonomic categories, we used them in the body-size based size class. Despite these attempts to achieve comparability, the food-web models likely lacked specific meio- and macrobenthic taxa. Hence, phyla rather than size classes should be considered when assessing the importance of polymetallic nodules for the food web.

Additionally, especially the food web developed for the Peru Basin missed specific phyla, such as meiobenthic Foraminifera (see Table S3) as there were no publications on this phylum.

However, this lack of literature was likely caused by a lack in specific taxonomic expertise during previous research cruises to the Peru Basin rather than caused by a deliberate omission of specific phyla. In contrast, the CCZ food web was based on a more balanced dataset for the different taxa due to the extensive baseline studies that were conducted in the CCZ during the last decade.

Authors reported some faunal groups only in a very coarse taxonomic resolution (see Table S3) which made it difficult to specify diets or feeding preferences. As a result, these compartments had a high number of trophic links which reduces the probability that a compartment is lost due to the loss of trophic interactions. Therefore, this study likely underestimated the importance of polymetallic nodules for food-web integrity, in particular for compartments of coarse taxonomic resolution or for compartments for which information about diet is poor.

The non-trophic interaction matrix only included commensalism and no other types of non-trophic interactions, such as competition (Kéfi et al., 2012). Interspecific competition for food resources might be one reason for sympatric speciation in abyssal deposit-feeding sea cucumbers: They have a high morphological diversity of tentacles that enabled the sympatric species to develop different feeding strategies and in this way overcome competition for food (Roberts & Moore, 1997). Hence, on evolutionary time scales, including competition in the food web would likely result in an increase in diversity and therefore an increase in compartments and links. In contrast, on the short term, competition would probably have no effect on the food-web structure and the non-trophic interactions that have been implemented in the model account for most of the secondary extinctions and shifts in food-web structure that can be expected from the absence of nodules.

### 4.2 Role of polymetallic nodules and stalked sponges in abyssal plain food webs

Our results consistently show for two polymetallic nodule-rich abyssal plains, the Peru Basin and the CCZ, which are ~6,000 km apart (Anonymous, 2018), that nodule removal will have knock-down effects on food-web integrity as it will result in the loss of food-web compartments due to the disturbance of trophic and non-trophic interaction webs. The loss of food-web compartments was almost exclusively the result of cascading effects from existing non-trophic interactions between nodules and sessile organisms and between sessile organisms attached to nodules and their associated fauna. Here, we discuss the importance of hard substrate for soft-sediment communities and we investigate the role of structural species versus highly-connected species in food webs. We also address potential consequences of future deep-seabed mining.

Hard substrate in soft-sediment communities increases habitat heterogeneity and may enhance species diversity and density (L. Buhl-Mortensen et al., 2012; Hasemann et al., 2013; Simon-Lledó et al., 2019; Vanreusel et al., 2016). In abyssal plains, this hard substrate is scarce and consists of clinker, hard-rock patches, polymetallic nodules, and glacial dropstones (Gooday et al., 2015; Hein, 2016; Kidd et al., 1981; Riehl et al., 2020). Dropstones may serve as island habitats (Ziegler et al., 2017) and host a diverse epifaunal assemblage of mainly Porifera, Chordata (in particular Tunicata), Echinodermata, Cnidaria, and Foraminifera (Gooday et al., 2015; Ziegler et al., 2017). At the West Antarctic Peninsula margin, for instance, dropstones contribute 20% to the total megabenthic species richness, though they cover less than 1% of the investigated seafloor (Ziegler et al., 2017). They may also increase epifaunal density, for instance in the Ardencaple Canyon in the Greenland Sea, where epifaunal density is positively correlated with the size of hard substrate (M. Schulz et al., 2010). In fact, dropstones can even modify hydrodynamics that lead to increased food supply to parts of the surrounding sediment and affect trophic structure, diversity, and life-history traits of nematodes (Hasemann et al., 2013).

Polymetallic nodules, in comparison, provide hard substrate for Arthropoda (in particular Crustacea), Bryozoa, Chordata (i.e., Tunicata), Cnidaria, Echinodermata, Foraminifera, Mollusca, Platyhelminthes, and Porifera (this study). Here, we determine that nodules are quantitatively important for filter and suspension feeders because 11% (Peru Basin) to 51% (CCZ) of all fauna are facultatively or obligatorily associated with the nodules. This close association may explain the positive correlation between nodule coverage and increased faunal density in the CCZ (Simon-Lledó et al., 2020). Furthermore, nodules and in particular stalks of attached epifauna can trap jelly-falls (Stratmann & van Oevelen, personal observations) and in this way increase the food supply for abyssal scavengers and omnivores. Hence, we conclude, that polymetallic nodules provide an important hard substrate in abyssal plains of the Pacific Ocean.

Our highly-resolved interaction food webs allowed to identify the species that have the so-called ‘highest impact’ (van der Zee et al., 2016). Surprisingly, at both sites, stalked-sponge species (i.e., *Hyalonema* sp. and *Caulophacus* sp.) were identified as ‘highest impact taxa’. Their removal had the largest impact on network properties because these sponges host commensal epibiota, such as filter and suspension feeders (e.g., Bivalvia, Crinoidea, Anthozoa, Cirripedia, and Crustacea), scavengers (e.g., Amphipoda, Mysida, and Isopoda), and predators (e.g., Polychaeta, Anthozoa, Ophiuroidea, and Isopoda). Sponge stalks were identified previously as ‘habitat islands’ for deep-sea fauna (Beaulieu, 2001a; Ilan et al., 1994) as they allow suspension feeders to move higher up into the benthic boundary layer, where higher flow velocities prevail (Lene Buhl-Mortensen et al., 2010). Hence, stalked sponges essentially extend the habitat of epifauna from the seafloor into the water column and are therefore considered ‘structural species’ (*sensu* (Huston, 1994)). An analysis of photograph transects at the abyssal Station M in the North-East Pacific showed that 87% of the stalks that emerged from the soft sediment were from the hexactinellid *Hyalonema* sp., of which only 14% were alive (Beaulieu, 2001b). These stalks host a diverse suspension feeding epibiota: The examination of 35 stalk communities showed that they harbored 8,580 individuals that could be classified into 139 taxa belonging to 13 different phyla (Beaulieu, 2001b). In the CCZ (this study) the stalks of *Hyalonema* sp. hosted 41 taxa from 4 different phyla and the stalks of *Caulophacus* sp. in the Peru Basin (this study) harbored 25 taxa belonging to 4 different phyla.

Theoretical food-web studies indicate that removal of highly connected species can lead to a cascade of secondary extinctions (Dunne et al., 2002; Sole & Montoya, 2001). In particular food webs with a low connectance (C ≤0.06) are sensitive to the loss of the most connected species (Dunne et al., 2002). Hence, we would expect that the removal of taxa with most trophic and non-trophic interactions in the CCZ (taxon with most trophic interactions: *Hymenaster* sp.; taxon with most non-trophic interactions: *Abyssarya* sp.) and the Peru Basin (taxon with most trophic interactions: *Actinia* sp.; taxon with most non-trophic interactions: *Munidposis* sp.) would lead to a cascade of secondary extinctions, because both webs have a low connectance (CCZ: C = 0.06, Peru Basin: C = 0.07; Table 1). Surprisingly, the removal of these most connected taxa did not cause any secondary extinctions. In other detritus-based food webs, removing the most connected species similarly had no to very little effect (Dunne et al., 2002; Warren, 1989; van der Zee et al., 2016) on the number of species and connectance, whereas the removal of the basal node, the detritus, resulted in a (rather trivial) immediate food-web collapse (Dunne et al., 2002). The CCZ- and the Peru Basin-food webs are likewise detritus-based, dominated by invertebrates, and the most connected species are generalist-feeding invertebrate predators (i.e., *Hymenaster* sp., *Actinia* sp.). Removing polymetallic nodules as primary habitat modifiers resulted in an increase in connectance by 2% (CCZ) to 23% (Peru Basin). This stresses the importance of non-trophic interactions in these nodule-rich abyssal plain food web, where the most connected species seem to be less important for overall food-web integrity because most can be considered generalist invertebrates. Therefore, we conclude, that in polymetallic nodule-rich abyssal areas in the Pacific, structural taxa contribute more to food-web integrity and species richness than highly connected taxa.

Our comprehensive analysis evidently and quantitatively shows the importance of polymetallic nodules and attached stalked sponges for the abyssal food web. It is therefore clear that the removal of polymetallic nodules due to deep-seabed mining would lead to a loss of food-web integrity and a significant depreciation of faunal biodiversity. It is not clear, whether this would affect the mined areas only or would result in regional or even global loss of biodiversity (Van Dover et al., 2017; Niner et al., 2018; Washburn et al., 2019). Ophiuroidea, for instance, have an unexpectedly high biodiversity in the CCZ (Christodoulou et al., 2019, 2020), and further discoveries of faunal biodiversity hubs in the poorly-studied deep sea (Ramírez-Llodrà et al., 2010) can be expected. In fact, we even neglected two very diverse taxonomic kingdoms, i.e., Bacteria and Archaea, in our study because we were not able to incorporate them adequately due to a poor understanding of their metabolic pathways. Hence, the polymetallic nodule-mining induced loss of biodiversity may be even higher than shown in this study.

Previous modelling studies about the impact of deep-seabed mining on the benthic ecosystem focused on carbon flows (de Jonge et al., 2020; Stratmann et al., 2018). These food-web models, however, group individual taxa in feeding types (van Oevelen et al., 2010; Soetaert & van Oevelen, 2009) which can mask biodiversity loss. Furthermore, the models concentrated on trophic interactions. However, for coastal ecosystems it is known that non-trophic interactions are important structuring factors in food webs (van der Zee et al., 2016). Therefore, we developed binary interaction webs including both trophic and non-trophic interactions. These webs show that non-trophic interactions are important for food-web integrity: The absence of polymetallic nodules will result in a reduced food-web complexity and a loss in biodiversity. Additionally, we identified stalked sponges as ‘structural species’ that host a high diversity of commensal fauna.

## 5 Conflict of Interest

The authors declare no conflict of interest.

## 6 Author Contributions

TS conceived the idea, searched the literature, compiled the datasets, developed the models, analyzed the results, and wrote the manuscript. DvO acquired funding, conceived the idea, developed the models, analyzed the results, and wrote the manuscript. DK identified sponges and its associated community on seafloor images, revised and edited the manuscript. KS acquired funding, revised and edited the manuscript.

## 7 Funding

The research leading to these results has received funding from the European Union Seventh Framework Programme (FP7/2007-2013) under the MIDAS project, grant agreement no. 603418, from the JPI Oceans – Ecological Aspects of Deep Sea Mining project (NWO-ALW grant 856.14.002), from the JPI Oceans – Impacts of deep-sea nodule mining project “MiningImpact 2” (NWO-ALW grant 856.18.003) and from the Bundesministerium für Bildung und Forschung (BMBF) grant n° 03F0707A-G. DvO and TS were also supported by the Dutch Research Council (DvO: NWO-VIDI grant no. 864.13.007, TS: NWO-Rubicon grant no. 019.182EN.012).

## 8 Acknowledgements

We thank chief scientists Pedro Martínez Arbizu (SO239) and Antje Boetius (SO242-2) and the captain and crew of RV Sonne for their excellent support during the SO239 ‘EcoResponse’ cruise and the SO242-2 ‘DISCOL revisited’ cruise. We further thank the ‘ROV KIEL 6000’ team from Geomar, Kiel (Germany).

## 10 Data Availability Statement

The datasets underlying the trophic and non-trophic interaction webs and the model code are attached as supporting information (Datasets S1 and S2, Programming file S1 and S2).

## Notes

### Competing Interest Statement

The authors have declared no competing interest.

